# Eta-secretase-like processing of the amyloid precursor protein (APP) by RHBDL4

**DOI:** 10.1101/2023.09.04.555897

**Authors:** Penalva Ylauna Christine Megane, Sandra Paschkowsky, Sherilyn Junelle Recinto, Anthony Duchesne, Clemence Levet, François Charron, Matthew Freeman, R. Anne McKinney, Jean-Francois Trempe, Lisa Marie Munter

**Author notes:** Correspondence should be addressed to Dr. Lisa Munter, Department of Pharmacology and Therapeutics, McGill University, Bellini Life Sciences Complex, Room 165, 3649 Promenade Sir William Osler, Montreal, QC, Canada H3G 0B1. These authors contributed equally (Penalva Ylauna Christine Megane, Sandra Paschkowsky, Sherilyn Junelle Recinto).

## Abstract

The amyloid precursor protein (APP) has been extensively studied with regards to its contribution to the pathology of Alzheimer’s disease. APP is an ubiquitously expressed type I transmembrane protein synthesized in the endoplasmic reticulum (ER) and translocated to the plasma membrane where it undergoes proteolytic cleavages by several identified proteases. Conversely to other known proteases, we previously elucidated human rhomboid protease RHBDL4 as a novel APP processing enzyme where several cleavages likely occur already in the ER. Interestingly, the pattern of RHBDL4-derived large APP C-terminal fragments resemble those generated by the η-secretase or MT5-MMP, which was described to generate so called Aη fragments. The similarity in large APP C-terminal fragments between both proteases raised the question whether RHBDL4 may contribute to η-secretase activity and Aη-like fragments. Here, we identified two cleavage sites of RHBDL4 in APP by mass spectrometry, which, intriguingly, lie in close proximity to the cleavage sites of MT5-MMP. Indeed, we observed that RHBDL4 generates Aη-like fragments *in vitro* without contributions of α-, β-, or γ-secretases. Such Aη-like fragments are likely generated in the ER since RHBDL4-derived APP-C-terminal fragments do not reach the cell surface. Inherited, familial APP mutations appear to not affect this processing pathway. In RHBDL4 knockout mice, we observed increased cerebral full length APP levels in comparison to WT brains in support of RHBDL4 being a physiologically relevant protease for APP. Furthermore, we found secreted Aη fragments in dissociated mixed cortical cultures from wild type mice, however significantly less Aη fragments in cultures from RHBDL4 knockout mice. Our data underscores that RHBDL4 contributes to η-secretease-like processing of APP and that RHBDL4 is a physiologically relevant protease for APP.

## Introduction

Alzheimer’s disease is the most common type of dementia, affecting 35 million people worldwide. It is a complex disease with multiple factors involved, including changes in vascular blood flow, inflammation, glucose metabolism, and cholesterol homeostasis (Duran-Aniotz & Hetz, 2016; Iturria-Medina *et al*, 2016; Puglielli *et al*, 2003; Spillantini & Goedert, 2013; Winblad *et al*, 2016). One pathological hallmark of Alzheimer’s disease is the production and aggregation of amyloid-β (Aβ) peptides leading to the characteristic amyloid plaques (O’Brien & Wong, 2011; Serrano-Pozo *et al*, 2011). Aβ peptides are generated through proteolytic cleavages from the larger amyloid precursor protein (APP). APP is an ubiquitously expressed type I transmembrane protein and of central importance in Alzheimer’s disease for the following reasons: 1) inherited mutations in APP cause a dominant, early onset form of familial Alzheimer’s disease (FAD) with full penetrance (Wu *et al*, 2012), 2) *APP* locus duplication is sufficient to inherit Alzheimer’s disease (McNaughton *et al*, 2012; Rovelet-Lecrux *et al*, 2006; Sleegers *et al*, 2006); and 3) genome-wide association studies (GWAS) revealed *APP* gene variants (single nucleotide polymorphisms) associate with an increased Alzheimer’s disease risk (Bellenguez *et al*, 2022; de Rojas *et al*, 2021; Schwartzentruber *et al*, 2021). Thus, genetic evidence showcases APP as a common denominator causatively linked to Alzheimer’s disease.

The exact physiological function of APP remains incomplete, but APP has been associated to pathways including cell-matrix and cell-cell interactions, synaptogenesis and axonal outgrowth (Anliker & Muller, 2006; Muller *et al*, 2017). Thus, a better understanding of APP’s biology may improve our understanding of the molecular processes involved in Alzheimer’s disease.

What has garnered interest in the field is the observation that APP undergoes proteolytic cleavages by many proteases, most of which show catalytic activity at the plasma membrane or the endo-lysosomal compartment leading to diverse soluble extra- or intracellular, and membrane-bound fragments (Andrew *et al*, 2016). Most prominent is APP processing by β- and γ-secretase yielding Aβ peptides (amyloidogenic processing), while APP processing by α- and γ- secretase prevents Aβ formation (non-amyloidogenic processing). Considering that Aβ species (of about ∼4 kDa) are hallmarks of AD pathology, such pathways have been well-characterized. Intriguingly, an earlier finding by Willem et al., sparked interest in a novel ∼8-14 kDa APP fragment subsequently named Aη (Willem *et al*, 2015). To generate Aη, APP undergoes a first cleavage by the matrix metalloproteinase-24 (MMP24), also known as membrane-type matrix metalloproteinase 5 (MT5-MMP) about 92 amino acids N terminal from the β-secretase cleavage site (Ahmad *et al*, 2006). Following MT5-MMP cleavage, α- or β-secretase would cleave the membrane bound C-terminal fragment and generate Aη fragments (Aη-α or Aη-β, respectively), and MT5-MMP was accordingly named η-secretase (Willem *et al*., 2015). Aη peptides impair long-term potentiation and strikingly, are about five times more abundant than Aβ peptides in human brain (Mensch *et al*, 2021; Willem *et al*., 2015). An Alzheimer’s disease mouse model (5xFAD) deficient for MT5-MMP showed reduced Aβ formation and maintained learning by promoting enhanced trafficking of APP to the endo-lysosomal compartment (Baranger *et al*, 2016a; Baranger *et al*, 2016b; Garcia-Gonzalez *et al*, 2021). Importantly, MT5-MMP-deficient mice alone showed reduced Aη formation, but notably some Aη was still formed, leading to the conclusion that while MT5-MMP contributes to the generation of Aη, other proteases may be involved in the formation of Aη fragments as well (Willem *et al*., 2015).

Independently, we posited the rhomboid protease RHBDL4 as an APP-processing enzyme *in vitro* (Paschkowsky *et al*, 2016). A striking characteristic of the RHBDL4-mediated APP cleavage is the emergence of at least six different, large C-terminal fragments (CTFs) of varying lengths between 10-15 and 20-25 kDa (Paschkowsky *et al*., 2016; Recinto *et al*, 2018). RHBDL4 resides in the endoplasmic reticulum (ER) (Fleig *et al*, 2012) and has the ability to cleave substrates not only in their transmembrane sequences but also in their ectodomains, thereby acting as both sheddase and intramembrane protease (Strisovsky *et al*, 2009; Wang *et al*, 2006; Wu *et al*, 2006). Considering its subcellular localization, we proposed that RHBDL4 cleaves APP in the ER in contrast to most other described APP proteases, i.e., α-, β-, and γ-secretase, meprin-β and MT5-MMP (Manucat-Tan *et al*, 2019). Thus, RHBDL4 catalytic activity on APP may modulate the amount of APP reaching the plasma membrane (Paschkowsky *et al*., 2016). Here, we identified two RHBDL4 cleavage sites within APP that lie in close proximity to the η- secretase cleavage site. We found that RHBDL4 alone is capable of generating Aη-like species without the activity of α- or β-secretase, which is further supported since large RHBDL4-derived APP CTFs do not reach the cell surface. Dissociated mixed cortical cultures from RHBDL4 knockout mice produced modest levels of Aη in comparison to wild type mice, further highlighting a role of RHBDL4 in APP biology.

## Results

### Identification of two RHBDL4 cleavage sites in APP

We previously demonstrated that cleavage of APP by RHBDL4 generates several novel, distinctive APP N- and C-terminal fragments (Paschkowsky *et al*., 2016). To identify the exact cleavage sites, we co-expressed myc-APP with either RHBDL4 or a catalytic inactive RHBDL4 S144A mutant in HEK293T cells. N-terminal APP fragments were immunoprecipitated from the supernatant using anti-myc-sepharose beads. Precipitates were digested with endoproteinase LysC and analyzed by mass spectrometry. Results were screened for APP peptides with a C-terminus not derived through LysC. If such unusual, truncated peptides were observed only with active, but not with inactive RHBDL4, we defined them as RHBDL4-cleavage sites. We identified a 20 amino acid long peptide corresponding to APP residues 494-526 (APP695 numbering) representing the full-length LysC-derived APP peptide in both samples (Fig. 1A, suppl. Fig. 1). In addition, two shorter, C-terminally truncated peptides were only identified in samples with active RHBDL4 (Fig. 1A). The two truncated LysC peptides demonstrate RHBDL4-specific cleavage sites at amino acids APP 505 (…ANM_505_-ISE…) and 514 (…SYG_514_-NDA…). Interestingly, these two cleavage sites lie just 1 and 10 amino acids C-terminal to the MT5-MMP (η-secretase) cleavage site at asparagine residue 504 (…AN_504_- MISE…), respectively (Fig. 1B).

**Figure 1:**
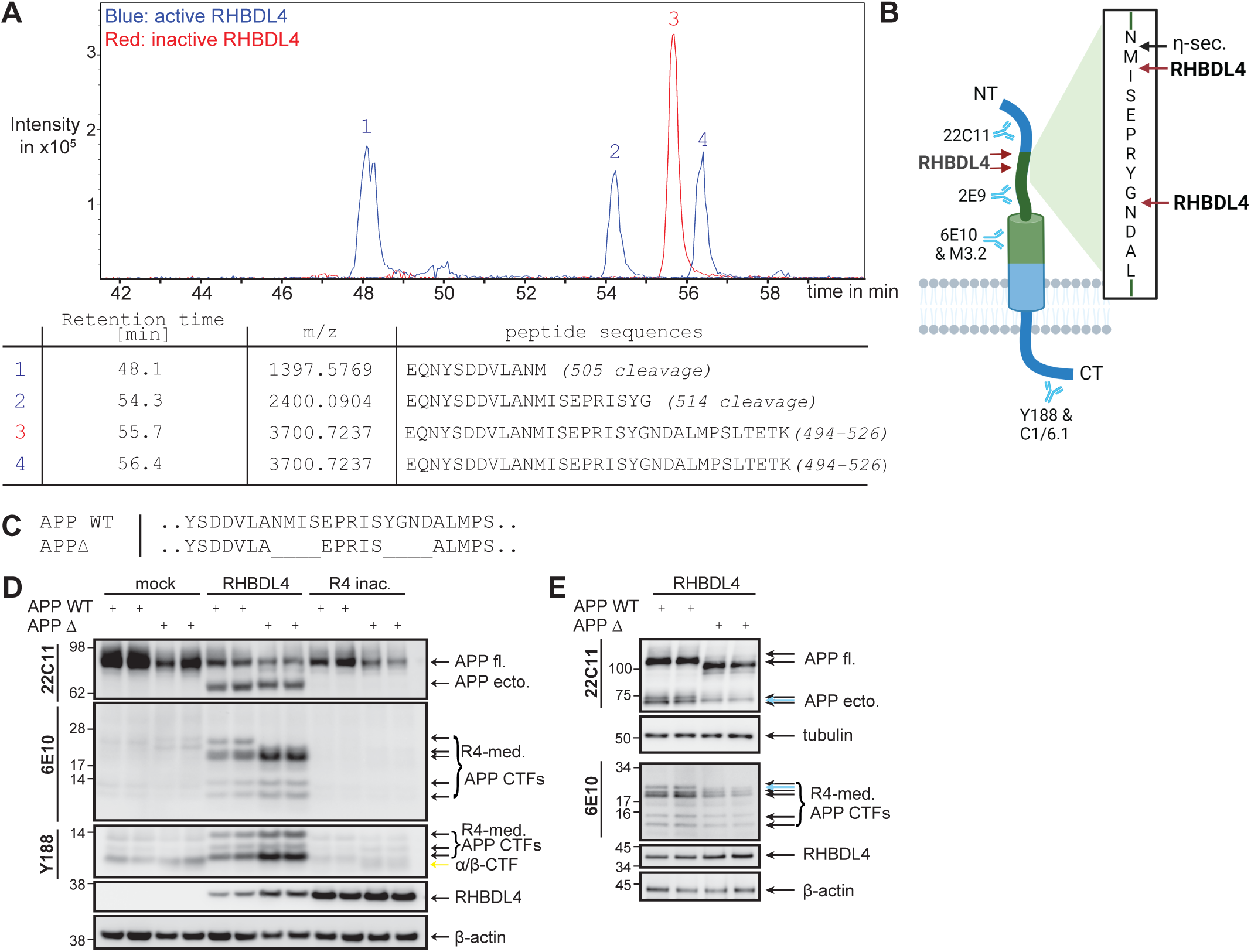
Identification of RHBDL4 cleavage sites in APP. **A)** Identification of RHBDL4 cleavage sites by mass spectrometry. Immunoprecipitation of N-terminally myc-tagged APP fragments after co-transfection with either active (ac.) or inactive (inac.) RHBDL4 in HEK293T cells. Samples were digested with LysC and analyzed by electrospray ionisation mass spectrometry (ESI-MS). Representative extracted ion chromatograms showing retention times for different identified peptides. The table lists the identified retention time per peak, the peptide mass per charge (m/z) and peptide sequences along with APP95 amino acid numbering for fragments or cleavage sites. Due to subtle differences in the automated injections, the retention time of the complete LysC peptide differs between the samples containing inactive RHBDL4 (55.7 min) and active RHBDL4 (56.4 min). **B)** Schematic representation of the identified RHBDL4 cleavage sites in APP created with BioRender.com. Antibody binding sites for 6E10, M3.2, 2E9, 22C11, Y188 and C1/6.1 antibodies are indicated. The previously identified η-secretase cleavage sites are indicated, scheme is not to scale. RHBDL4 has at least 4 more cleavage sites further towards the APP C-terminus which are not indicated (see Fig. 5 or (Paschkowsky *et al*., 2016). **C-E)** Analysis of RHBDL4-mediated (R4-med.) cleavage of the APP deletion (APPΔ) mutant. Amino acid stretches comprising two amino acids N- and C-terminal of both identified cleavage sites were deleted (as shown in C). Comparison of RHBDL4 cleavage pattern for APP WT and APPΔ upon co-transfection. Different gel systems were used to optimally analyse the fragments, 4-12% bis-tris (D), 8% tris-glycine (upper panel E) and 10-20% tris-tricine (lower panel E). Blue arrows indicate novel bands in the APPΔ samples. Detection of APP full length (APP fl.) and APP ectodomain (APP ecto.) with 22C11, CTFs with 6E10 and Y188; RHBDL4 with anti-myc antibody. β-actin or tubulin as loading controls. Representative western blot of 3 individual experiments is shown.

To validate the cleavage sites, we introduced mutations in APP to prevent RHBDL4 cleavage. Sequence recognition motifs of rhomboid proteases have been proposed for transmembrane sequences (Strisovsky *et al*., 2009). In fact, single point mutations have been ineffective at abrogating cleavages. Therefore, we generated an APP deletion mutant (APP△), in which 4 residues comprising P2’-P2 at both cleavage sites were deleted (Fig. 1C). Co-expression of APP△ with RHBDL4 revealed the lack of the largest APP-CTF while all other CTFs appear similar when compared to APP wild type (WT) (Fig. 1D). To better resolve effects of APP△ on the cleavage fragments, samples were separated on 8% SDS-PAGE gels. This higher resolution indeed showed that one of the two 70-73 kDa corresponding APP ectodomain bands is missing in APP△ but not in APP WT, while the remaining band runs in between the two bands seen for APP WT (blue arrow) (Fig. 1E). Furthermore, using a 10-20% tris-tricine gel for the CTFs, we confirmed the lack of the largest CTF, but also observed a faint, additional band migrating at a different molecular weight as the CTFs from APP WT (Fig. 1E; blue arrow). The origin of this additional band has not been identified; it could derive from either an alternative RHBDL4 cleavage triggered by the amino acid deletions or from a different migration behaviour of a large CTF containing the deleted region(s). Overall, the results show two RHBDL4 cleavage sites responsible for yielding two major RHBDL4-derived APP fragments.

### APP FAD mutations do not affect RHBDL4-mediated processing of APP

To evaluate the role of RHBDL4-mediated APP processing in FAD, we assessed whether APP FAD mutations could affect RHBDL4 cleavages as they do for other APP processing pathways. We therefore co-expressed RHBDL4 with 8 different APP FAD mutations in HEK293T cells and quantified the formation of APP-CTFs (Cacace *et al*, 2016). For the 20-25 kDa large, RHBDL4-derived APP-CTFs, we observed no significant changes for APP V715M, I716F and V717F mutants, however, a significant increase for the APP Swedish mutant as compared to APP WT (Fig. 2A, B). For the 10-15 kDa APP-CTFs, we observed no significant changes as compared to APP WT (Fig. 2A, B). Thus, the APP Swedish mutant may not only affect the processing efficiency of β-secretase, but also that of RHBDL4, while the effect of other APP FAD mutants on RHBDL4-mediated APP processing is negligible

**Figure 2:**
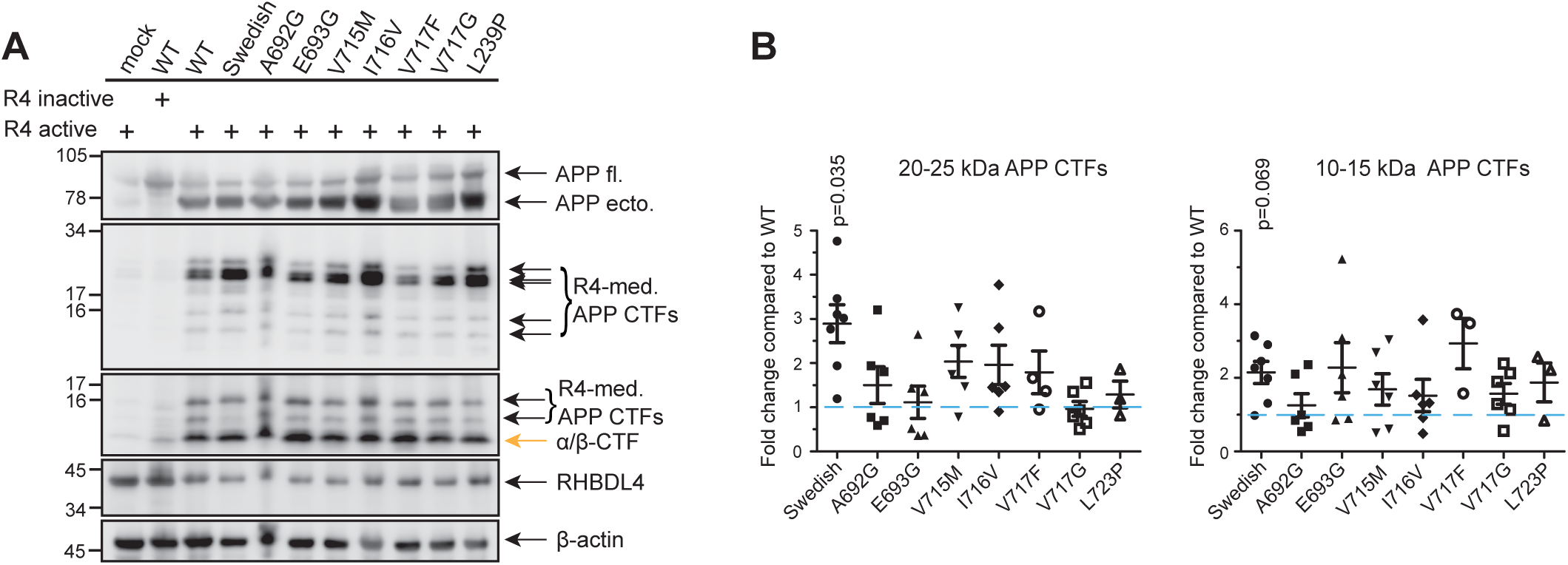
Familial APP mutations do not affect RHBDL4-mediated processing of APP. **A-B)** RHBDL4-mediated (R4-med.) processing of familial AD mutants of APP. Cotransfection of various familial APP mutants with active RHBDL4. Detection of APP full length (APP fl.) and APP ectodomain (APP ecto.) with 22C11, CTFs with 6E10 and Y188; RHBDL4 with anti- myc antibody. β-actin as loading controls. APP-CTFs were quantified and normalized first to β-actin and then the fold change compared to WT was calculated. WT was always set to 1 in each individual experiment (blue dashed line). Mean ± SEM is displayed, n=3-6, p-values for Bonferroni-corrected one sample t-tests are reported.

### RHBDL4-mediated large APP-CTFs are not translocated to the plasma membrane

The close proximity of the RHBDL4-cleavage sites to the MT5-MMP cleavage site (Ahmad *et al*., 2006), as well as the similarity of APP-CTF patterns deriving from MT5-MMP (Willem *et al*., 2015) or RHBDL4 (Paschkowsky *et al*., 2016) processing, raised the question whether RHBDL4 could contribute to the formation of Aη-like fragments. For this to occur, the larger RHBDL4-derived APP-CTFs would need to traffic to the cell surface or the endosomes to allow either α-, or β-secretase to generate the C terminus of Aη fragments (Kinoshita *et al*, 2003; Willem *et al*., 2015). Therefore, we evaluated the cell surface presence of RHBDL4-derived large APP-CTFs by cell surface biotinylation in cells overexpressing APP with either active RHBDL4, catalytically inactive RHBDL4, or a different rhomboid protease, RHBDL2 as negative control. We observed large APP-CTFs abundantly present in total cell lysates, albeit no large APP-CTFs were detected at the cell surface (Fig. 3A). Consistently, full-length APP was reduced when co-expressed with active RHBDL4, which degrades it in the ER, in contrast to when inactive RHBDL4 or RHBDL2 were co-expressed, which allow full length APP to traffic to the cell surface. Collectively, these findings demonstrate that RHBDL4-mediated large APP-CTFs that are not trafficked to the cell surface and thus are probably not further processed by α- or β-secretases. The results also imply that RHBDL4-mediated APP processing in the ER may impact APP physiology by regulating its plasma membrane presence.

**Figure 3:**
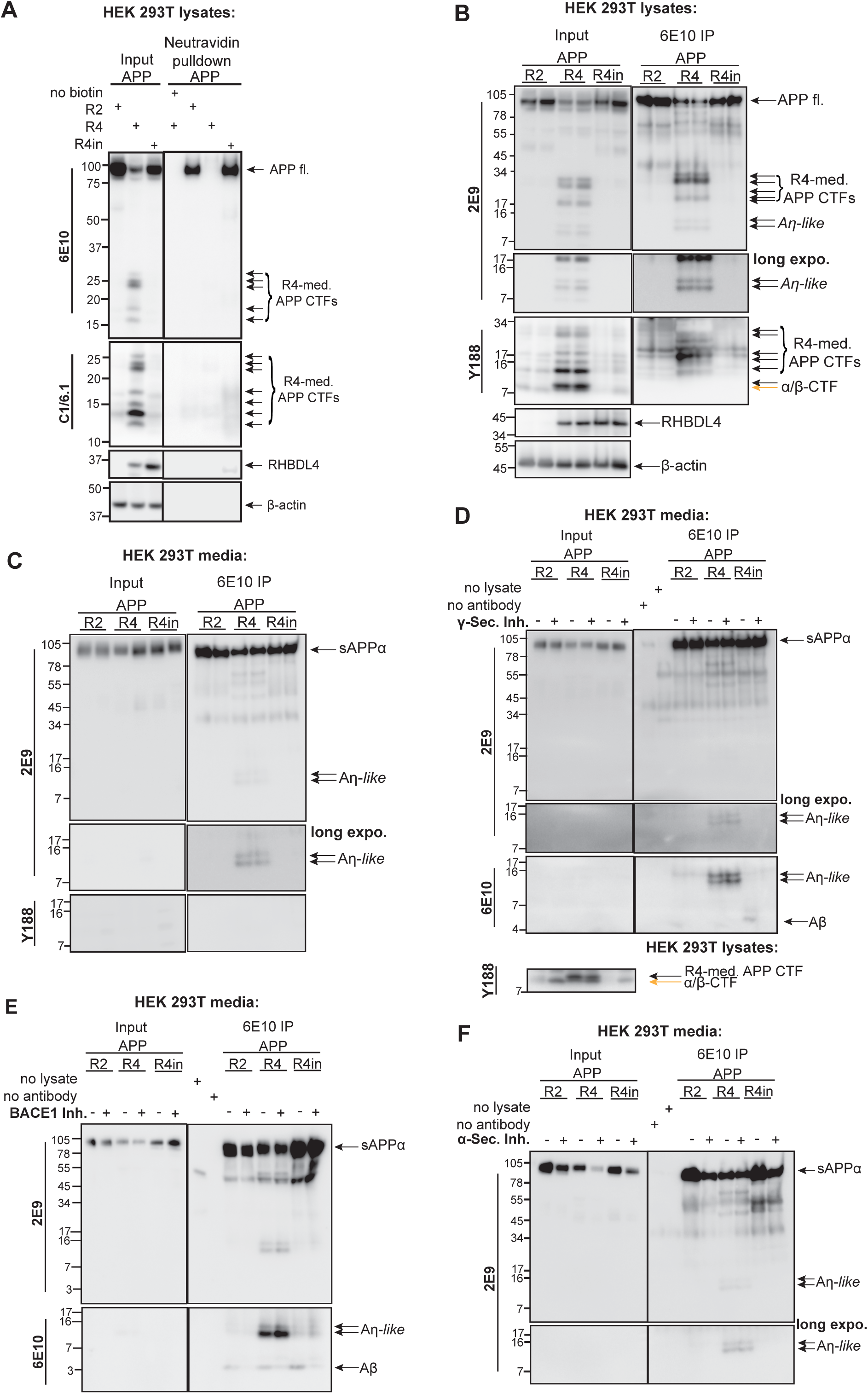
RHBDL4 generates Aη-like peptides *in vitro*. **A)** Investigation of RHBDL4 mediated APP CTFs at the cell surface using cell surface biotinylation. Co-transfection of APP and RHBDL2 (R2), active RHBDL4 (R4) or inactive RHBDL4 (R4in). The input consists of lysates without neutravidin to serve as loading controls (left panels). Biotinylated cell surface proteins were pulled down using neutravidin (right panels). Detection of APP fl. with 6E10, CTFs with 6E10 and C1/6.1; RHBDL4 with rabbit anti- RHBDL4 antibody. Representative western blot of 3 individual experiments is shown. **B-C)** Immunoprecipitation of Aη species from cell culture supernatant and lysate. Co-transfection of APP and RHBDL2 (R2), active RHBDL4 (R4) or inactive RHBDL4 (R4in). Total cell culture lysates or supernatant are used as input (left panels) while immunoprecipitation (IP) was performed using the 6E10 antibody (right panels). Detection of APP full length (fl.), sAPP11 and Aη with 2E9, CTFs with Y188; RHBDL4 with rabbit-anti-RHBDL4 antibody and β- actin as a loading control. Representative western blot of 3 individual experiments is shown. **D-F)** RHBDL4-mediated Aη generation is independent of canonical processing by 11-, β- or γ- secretases. Cells were treated with either 11-secretase inhibitor (11-Sec. Inh.), BACE-1 inhibitor (BACE1 Inh.) or γ-secretase inhibitor (γ-sec. inh). Total cell culture supernatant is used as input (left panels) while immunoprecipitation (IP) was performed using the 6E10 antibody (right panels). Detection of sAPP11 and Aη with 2E9 and 6E10. Representative western blots of each inhibitor experiments are shown, n=3 per inhibitor.

### RHBDL4 generates Aη-*like* peptides

RHBDL4-derived APP-CTFs do not reach the cell surface (Fig 3. A) and are not dependent on α- or β-secretase activity (Paschkowsky *et al*., 2016). However, since RHBDL4 has at least 4 other cleavage sites in APP located further C-terminal to the two identified cleavage sites (see schematic Fig. 5, Paschkowsky *et al*., 2016), we investigated if RHBDL4 directly could generate fragments similar to Aη, without the involvement of α- or β-secretase. Thus, we performed immunoprecipitation (IP) experiments from cell culture supernatants and lysates of HEK293T cells transfected with APP and either RHBDL4, inactive RHBDL4 or RHBDL2. IPs were performed with the 6E10 antibody binding in the Aβ region of APP; western blots were first stained with the 2E9 antibody (Fig. 3B). In cell lysates, we detected 2E9-immunoreactive bands at the anticipated molecular weight of 8-14 kDa, which were, however, difficult to distinguish from APP CTFs detected with an antibody binding to the very C terminal amino acids of APP (Y188), which is not present in Aη fragments (Fig. 3B). Since Aη peptides can be secreted, we also performed IPs on the cell culture supernatants. Immunoprecipitation of supernatants with 6E10 from cells overexpressing active RHBDL4 and APP elucidated double-band signals between 8 and 14 kDa that resemble the Aη fragment pattern previously published by Willem et al. (Fig. 3C) (Willem *et al*., 2015). To differentiate Aη fragments from those similar fragments derived by RHBDL4, we herein call these Aη-like fragments. These Aη-like fragments could not be detected in the cell supernatant when RHBDL2 or inactive RHBDL4 S144A were co-expressed. Further, Aη-like fragments were not immunoreactive with the antibody Y188 directed against the APP C-terminus (Fig. 3C).

To further evaluate the potential involvement of α-, β- or γ-secretase in Aη-like generation, we performed similar IP experiments in the presence of either α-, β- or γ-secretase inhibitors (Fig. 3D-F). None of the treatments impacted the RHBDL4-mediated generation of Aη-like fragments. As expected, we observed lower levels of sAPPα and Aβ in the cell supernatant upon α- or β- secretase inhibition, respectively (Fig. 3E, F), while increased levels of α/β-CTFs upon γ- secretase inhibition (Fig. 3D), indicative of effective treatments. These findings coincide with our prior report that RHBDL4-derived APP-CTFs are independent of the canonical APP processing pathways (Paschkowsky *et al*., 2016). Taken together, we propose that Aη-like fragments are derived entirely from RHBDL4-mediated APP cleavages at both N- and C-termini and thus are generated intracellularly in the ER, and subsequently, potentially passively, secreted.

### RHBDL4-mediated Aη-like fragments in the brain

To study the physiological relevance of RHBDL4-generated Aη-like fragments in the brain, we extracted Aη from brain tissues of homozygous RHBDL4 knockout (KO) and wild type mice as shown previously (Willem *et al*., 2015). However, in our hands, Aη detection by Western blot was frequently perturbed by an unidentified negative signal preventing sound interpretation of results (Fig. 4A). The Aη signals appeared similar between RHBDL4 KO and wild type mice which we suspect may be due to the cellular heterogeneity that exists in the brain and the expression of other known APP-cleaving enzymes that may also vary across cell types. Further, we quantified the expression of endogenous full-length APP from 10-11 months old mouse brain lysates by Western blot, and found about 1.5-fold higher APP expression in brains of RHBDL4 KO mice compared to wild type littermate controls (Fig. 4B, C). This finding implies that APP is a physiological relevant substrate of RHBDL4, so that upon loss of the protease the substrate accumulates. Henceforth, we turned to primary dissociated mixed cortical cell cultures to determine if Aη-like fragments could be generated by RHBDL4. Cultures from neonatal mice were cultivated for 14 days before Aη fragments were immunoprecipitated from the supernatants by the M3.2 antibody recognizing endogenous mouse APP (Fig. 1B, 4D). Of note, no difference in APP expression was observed between wild type and RHBDL4 KO primary cortical cultures lysates, while it was observed in total brain lysates (Fig. 4E, F). This can likely be attributed to the difference in age where 10-11 months old mice were used for the lysates, but P0-P1 pups for the dissociated cortical cultures. Strikingly, we observed almost 90% reduction in Aη production in mixed cortical cultures supernatants from RHBDL4 KO as compared to wild type mice (Fig. 4G, H). Moreover, treatment with metalloprotease inhibitors TIMP2 or BB94 in wild type and RHBDL4 knockout cultures further reduced Aη formation, confirming that both metalloproteases and RHBDL4 play a role in Aη production (Fig. 4G, H). Overall, we demonstrated that RHBDL4 contributes to Aη-like processing *ex vivo*.

**Figure 4:**
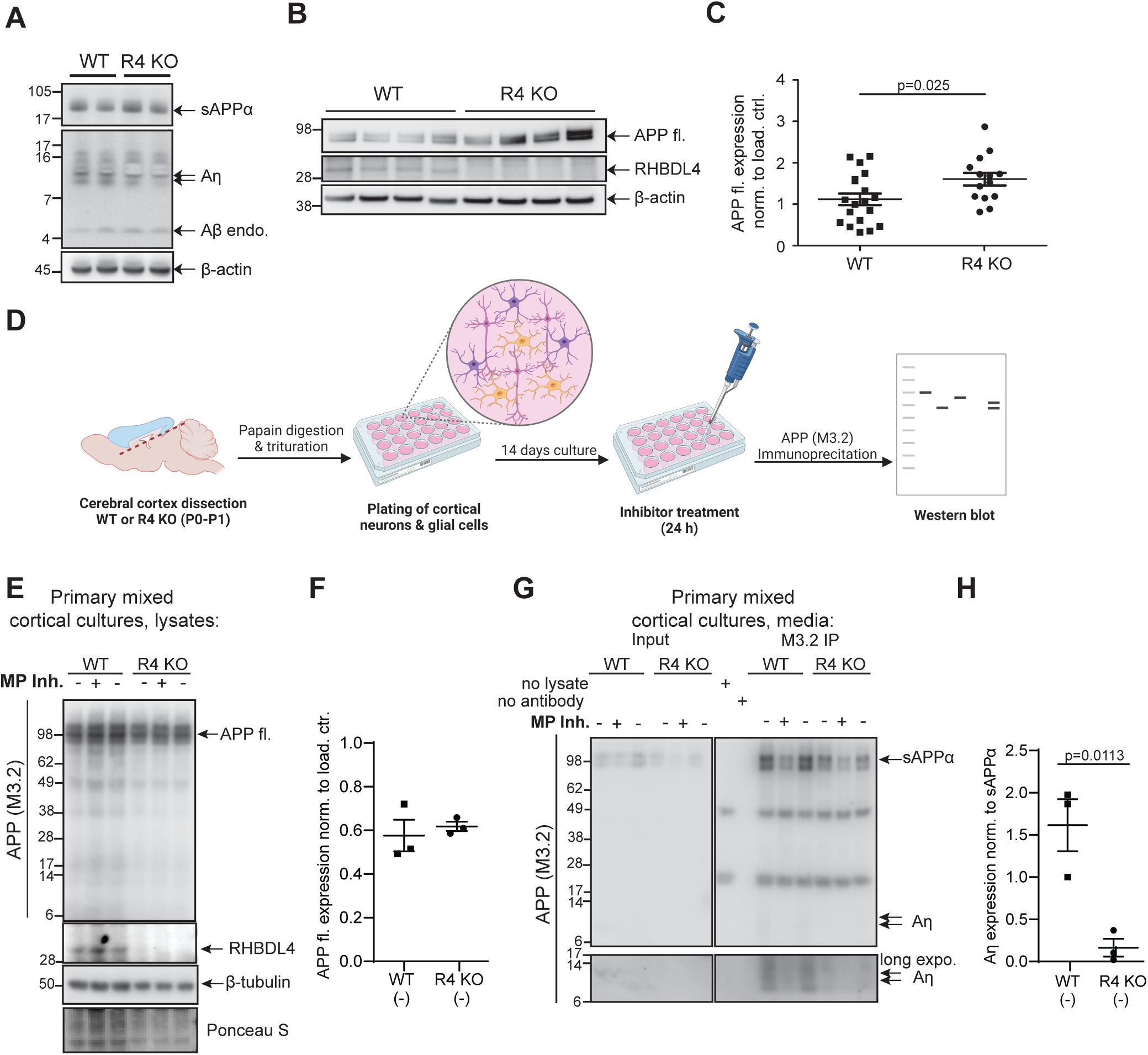
RHBDL4 knockout affects APP levels and Aη production *in vivo*. **A)** Aη extraction was performed according to (Willem *et al*, 2006), extraction of soluble proteins upon DEA extraction from brain tissue homogenates of WT and RHBDL4 KO mice at 10-11 months of age. sAPPα and Aη were detected with M3.2 antibody (specific for mouse Aβ), β-actin as loading control. Representative western blot for n=8 brain samples (WT and KO, each). **B-C)** Full length APP levels in brain tissue lysates of 10-11 months old RHBDL4 knockout (R4 KO) mice as compared to age matched wild type (WT) mice. Equal amounts of protein were loaded per lane. Detection of APP full length (fl) with 22C11, endogenous RHBDL4 with rabbit anti-RHBDL4 antibody, and β-actin as a loading control. Quantification with ImageJ, normalized to β-actin, mean ± SEM, n=14-19, p-value for unpaired two-tailed t-test is reported. **D)** Schematic representation of cortical dissociation and primary cell culture procedure followed by Aη immunoprecipitation from cell medium and downstream analysis via western blot. Created using BioRender. **E-F)** Full length APP expression from primary cortical cell culture lysates prepared from WT or RHBDL4 knockout brains. Representative western blot of 3 individual experiments is shown; APP quantification of untreated condition normalized to ponceau S with ImageJ, mean ± SEM, unpaired two-tailed t-test performed. Treatment with metalloprotease inhibitor (MP inh.) TIMP2. Detection of APP full length (fl.) with M3.2, endogenous RHBDL4 with rabbit-anti-RHBDL4 antibody, β-tubulin and ponceau S as loading controls **E-F)** Immunoprecipitation of Aη species from primary cortical cell culture supernatant prepared from WT or RHBDL4 knockout mouse brains. Input consists of total cell culture supernatant (left panels) while immunoprecipitation (IP) was performed using the M3.2 antibody (right panels). sAPP and Aη detection using M3.2 antibody. Representative western blot of 3 individual experiments is shown; quantification of untreated condition with ImageJ normalized to sAPP11, mean ± SEM, p-value for unpaired two-tailed t-test is reported. Treatment with metalloprotease inhibitor (MP inh.) TIMP2.

## Discussion

Although they were initially discovered as intramembrane proteases, rhomboid proteases are able to cleave substrates both in the plane of the membrane as well as in their ectodomains (Avci & Lemberg, 2015; Bock *et al*, 2022; Paschkowsky *et al*., 2016; Recinto *et al*., 2018; Strisovsky *et al*., 2009). Here, we demonstrate that RHBDL4-mediated cleavages of APP in the ER generate Aη-like peptides. Considering that those Aη-like peptides require neither α-, β- nor γ-secretase activity, our data suggests that RHBDL4 activity itself is sufficient for producing these fragments via multiple cleavage events in different positions to generate various N- and C-terminal fragments (Fig. 5). Interestingly, the two RHBDL4 cleavage sites we identified are in close proximity to the proposed η-secretase cleavage site (Ahmad *et al*., 2006; Willem *et al*., 2015). Please note that other RHBDL4 cleavage sites have yet to be characterized as previous findings indicate that RHBDL4 cleaves APP at least 6 times (Paschkowsky *et al*., 2016). We corroborated these cleavage sites by generating a deletion mutant which abrogates the formation of the largest APP-CTF. Of importance is the highly similar pattern of APP CTFs generated by RHBDL4 to those observed by Willem et al., who also identified about 5 APP CTFs ranging between approximately 10 – 30 kDa (Willem *et al*., 2015). Hence a single cleavage by MT5-MMP would not explain this complex CTF pattern, implying for the significance of RHBDL4’s η-secretase-like activity.

**Figure 5:**
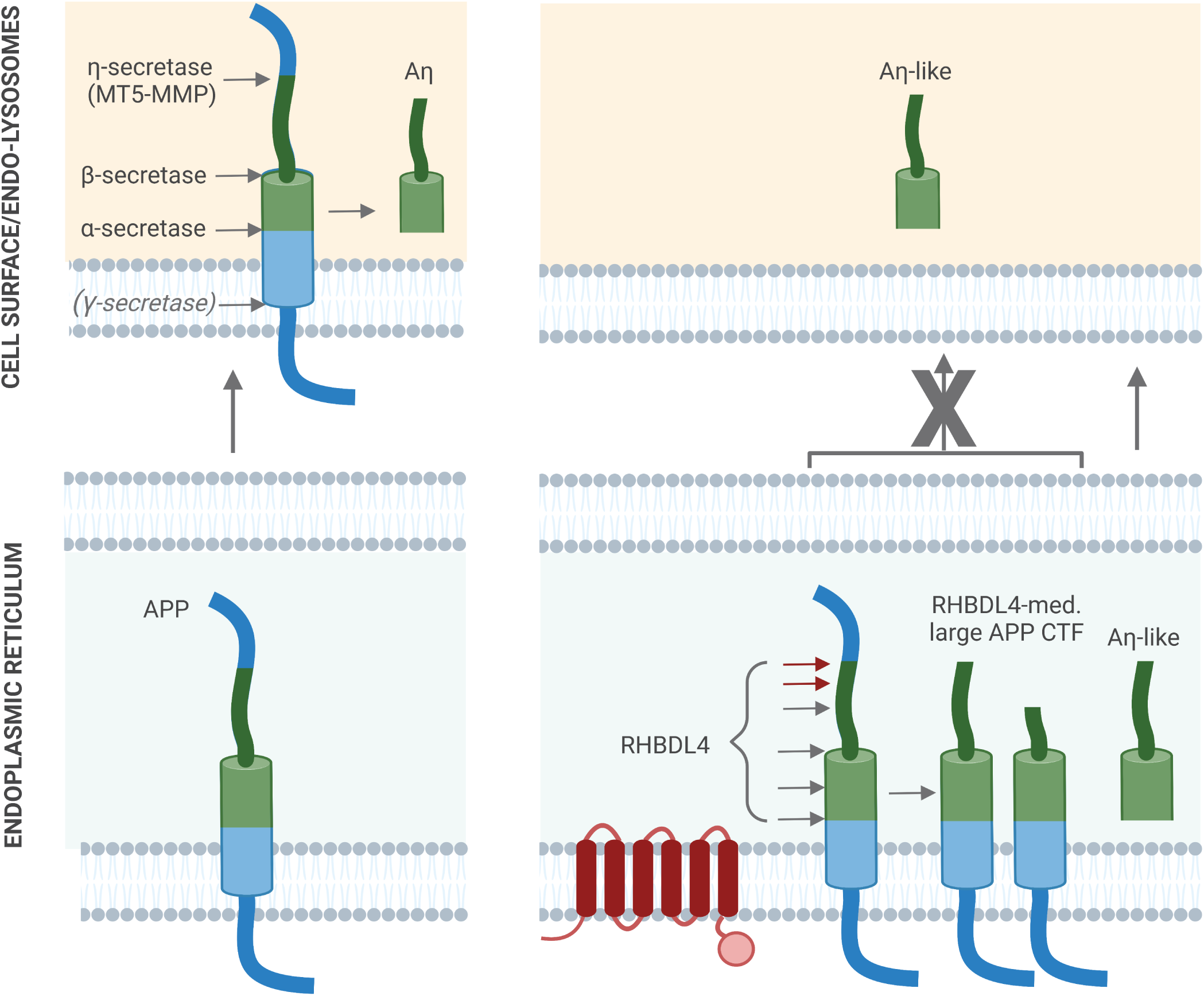
Scheme of APP processing and Aη formation in different compartments. In the absence of RHBDL4, full length APP traffics to the cell surface where MT5-MMP as well as α- or β-secretases will process APP to generate Aη at the cell surface (left panels). In the presence of RHBDL4, APP will be cleaved by RHBDL4 in the ER and RHBDL4-derived large APP C-terminal fragments do not reach the cell surface. RHBDL4-derived Aη-like peptides are directly generated in the ER (right panels).

The physiological relevance of the APP processing pathway leading to Aη peptides remains unclear. Due to the higher abundance of Aη peptides in the brain than the well-established Aβ peptides (Willem *et al*., 2015) and their potential involvement in impairing long-term potentiation (LTP) (Mensch *et al*., 2021), it is intriguing to speculate that this non-canonical APP processing pathway may be implicated in Alzheimer’s disease. It is therefore necessary to investigate the *in vivo* relevance of these pathways to understand their potential involvement in Alzheimer’s disease. A remarkable feature of APP is its processing by a myriad of proteases, besides α-, β- and γ-secretases, multiple others such as MT5-MMP, RHBDL4, HtrA2, meprin-β, and possibly MT3-MMP. However, it remains elusive which pathways are active in which cell types at what stage and how they may be regulated (Ahmad *et al*., 2006; Bien *et al*, 2012; Huttunen *et al*, 2007; Park *et al*, 2006; Zhang *et al*, 2015). The relevance of Aη peptides in Alzheimer’s disease is also unclear. Functional redundancy between rhomboid proteases and sheddases, especially metalloproteases, is not unprecedented. In particular, the human rhomboid protease RHBDL2 and ADAMs possess overlapping substrate specificity with B-type ephrins (Pascall & Brown, 2004) and EGF (Adrain *et al*, 2011; Johnson *et al*, 2017). Hereby, we propose that RHBDL4-mediated Aη generation is a physiological event in the ER, likely contributing to elevated extracellular Aη levels in the brain. Not only have we observed higher levels of the RHBDL4 substrate - full-length APP - in RHBDL4 KO compared to wild type mice, but we also found significantly lower Aη-like fragments in primary mixed cortical cultures in the absence of RHBDL4. In addition, RHBDL4 activity on APP in the ER could further serve as a mechanism to regulate the cell surface levels of full-length APP (Fig. 5). Taken together, our findings underscore that RHBDL4 is a physiologically relevant protease in APP biology.

The effect of MT5-MMP has previously been investigated in mouse models of Alzheimer’s disease, i.e., MT5-MMP knockout mice were crossed with the 5xFAD model of amyloidosis (Baranger *et al*., 2016a; Baranger *et al*., 2016b). The authors observed maintained cognitive performance and decreased formation of Aβ peptides indicating that MT5-MMP could serve as a new drug target for Alzheimer’s disease. Another research group found that MT5-MMP cleaves APP in response to oxidative stress further linking MT5-MMP to potentially pathogenic pathways in Alzheimer’s disease (Llorente *et al*, 2020). Mechanistically, the authors report that MT5-MMP appears to affect APP processing through proteolytic and non-proteolytic functions: MT5-MMP appears to promote the trafficking of APP towards the endolysosomal compartment which would lead to Aβ generation, a pathological hallmark of Alzheimer’s disease (Garcia-Gonzalez *et al*., 2021). Similarly, RHBDL4 was meanwhile reported to carry proteolytic as well as non-proteolytic functions. The Freeman group reported that RHBDL4 protects from ER stress by interacting with CLIMP-63, a protein shaping the morphology of the ER and stabilizing ER sheets (Lastun *et al*, 2022). Likewise, RHBDL4 was suggested to organize GPCR-mediated trafficking events like for the signaling molecule TGFα (Wunderle *et al*, 2016). Using BioID proximity assays, it was indeed shown that RHBDL4 interacts with dozens of other proteins likely through interactions not related to proteolysis further supporting non-catalytic functions of RHBDL4 (Hsiao *et al*, 2023; Ikeda & Freeman, 2019). Remarkably, we found an increased catalytic activity of RHBDL4 at low cellular cholesterol levels (Paschkowsky *et al*, 2018), while β- and γ-secretases show an increased activity at high cellular cholesterol levels (Holmes *et al*, 2012; Kalvodova *et al*, 2005; Marquer *et al*, 2011; Osenkowski *et al*, 2008). Interestingly, APP itself is a cholesterol-binding protein (Barrett *et al*, 2012; Pester *et al*, 2013; Pierrot *et al*, 2013). Thus, it is tempting to speculate that APP processing in the ER by RHBDL4 versus at the cell surface/endolysosomes by β- and γ-secretases (see Fig. 5) could be counter-regulated by cellular cholesterol levels and modulate the physiological function of APP. The production of Aη-like or Aβ fragments would then be, in part, markers of the respective degradation pathway.

## Conclusion

In summary, we aimed to shed light on the relevance of RHBDL4-mediated APP processing. We identified RHBDL4 as a physiological protease with significant relevance for APP biology, contributing to Aη-like peptide production. Further research will be necessary to understand the regulation and implication of RHBDL4-mediated proteolytic APP degradation in the ER.

## Materials and Methods

### DNA constructs

Plasmid pCMV6 encoding cDNA for human RHBDL2 and RHBDL4 with a C-terminal myc-FLAG tag was obtained from OriGene, USA. cDNA encoding APP695 untagged (in pcDNA3.1, Invitrogen) was a kind gift of Dr. Claus Pietrzik, Johannes Gutenberg University, Mainz, Germany. For mass spectrometry and cell culture experiments, APP695 with an N-terminal myc tag and a C-terminal FLAG Tag was cloned into pcDNA3.1. The APPΔ construct was designed using geneblock technology (IDT) with the following strategy: 4 ng/uL of geneblock was digested using high-fidelity BamHI and EcoRI restriction enzymes in a 25 µL reaction volume at 37°C overnight (16-18 h). Similarly, 1 µg of pcDNA3.1 containing APP695 cDNA plasmid was digested using the same restriction enzymes. After digestion, all samples were subjected to a calf intestinal alkaline phosphatase (CIP) treatment at 37°C, followed by heat inactivation at 65°C. 50 ng of gel-purified vector was ligated to the insert at RT overnight (16-18 h). The recombinant construct was transformed into competent cells via heat shock. Single bacterial colonies were picked and cultured overnight. Empty vectors pcDNA3.1 or pCMV6 were used as negative controls. Inactive RHBDL4 S144A was generated by site directed mutagenesis using the forward primer 5’-GCTGTAGGTTTCGCAGGAGTTTTGTTT-’3 and the reverse primer 5’-AAACAAAACTCCTGCGAAACCTACAGC-’3. All expression vectors were verified by dideoxy DNA sequencing (McGill Génome Québec Sequencing Center).

### *In vitro* cultures and transfection

HEK293T cells were cultivated in Dulbeccós modified Eagle medium (DMEM) containing 4.5 g/l glucose, 0.584 g/l L-glutamine and 0.11 g/l sodium pyruvate (Wisent), supplemented with 10% fetal calf serum (FCS) (Wisent), at 37°C and 5% CO_2_, and were regularly passaged at a confluency of 80-90%. 6 x 10^5^ cells per 6-well were seeded 24 h before transfection. Transient transfection was performed with 2 µg DNA and 4 µl polyethylenimine (PEI) per well. For co-transfection, a DNA ratio of APP to rhomboid protease of 5:1 was used. 36 h after transfection, cells were lysed with TNE-lysis buffer (50 mM Tris, pH 7.4, 150 mM NaCl, 2 mM EDTA, 1% NP40, and complete Protease Inhibitors, Roche) for 30 min on ice then spun down at 11,000 rpm for 10 min. Lysates are collected and prepared for SDS-polyacrylamide gel electrophoresis (SDS-PAGE).

### Mouse primary cortical cultures and treatments

*Ex vivo* experiments were performed on primary cortical cultures prepared from early postnatal C57BL/6 WT or RHBDL4-knockout mice as previously described (Brewer & Torricelli, 2007; Deane *et al*, 2013; Gao *et al*, 2019). In brief, post-natal day (P) 0–1 mice pups were decapitated, their brains removed, and the cortices microdissected. These cortices were maintained in chilled HBSS supplemented with 0.1LJM HEPES buffer and 0.6% glucose, then digested with 165 U papain for 20 min in a shaking water bath at 37°C. Neurons and glia were dissociated by trituration and suspended in DMEM supplemented with 1% penicillin-streptomycin, 10% FBS, and 0.6% glucose. Cells were then plated onto poly-d-lysine-coated 10 mm coverslips at an approximate density of 12,000 cells/cm^2^ and placed in an incubator at 37°C. 24 h later, plating media was replaced with Neurobasal-A growth media supplemented with 2% B-27 supplement, 1% GlutaMAX, and 1% penicillin-streptomycin. Cultures were then fed every 3–4 d and allowed to mature until 14 days in vitro (DIV) at 37°C in a humidified environment of 5% CO_2_. Mouse primary cortical cultures were then treated with 0.5 μg/ml human recombinant tissue inhibitor of metalloproteinase-2 (TIMP2, Sigma) for one biological replicate or 10 μM BB94 for two biological replicates, for 24 h.

### Cell surface biotinylation

36 h following transfection, cells were washed twice with pre-warmed PBS containing Mg^2+^ and Ca^2+^ ions (Wisent) and once with pre-chilled Mg^2+^/Ca^2+^ PBS. On ice, cells were incubated in 0.5 mg/mL non-permeant, EZ-Link Sulfo-NHS-SS-Biotin (Thermo Scientific) diluted in Mg^2+^/Ca^2+^ PBS for 10 min. Residual unbounded biotin was washed away twice with PBS containing 10 mM glycine (BioShop). Membrane extracts were lysed with TNE buffer and lysates were collected as described above. 50 μL total cell lysates were taken as expression controls and prepared with 1 x LDS sample buffer with 10% β-mercaptoethanol. Remaining lysates were diluted with PBS and incubated rotating overnight at 4°C with 60 μL washed, diluted neutravidin-beads (1:1 beads: PBS, Thermo Scientific). Samples were spun down at 1,500 x g for 3 min repeatedly until final wash. Beads were washed twice with 400 mM NaCl in PBS and once with PBS only. Using an 18-gauge needle, beads were dried. 2 x LDS sample buffer and additional 5% SDS were added then samples were boiled for 5 min at 65°C.

### Immunoprecipitation for Aη from supernatant and lysate

For HEK293T cells, 36 h after transfection, media was changed to DMEM plus 2% FCS and cells were conditioned for 24 h. Treatments with 10 µM α-secretase inhibitor [BB94 (Abcam)], or 1 µM β- or γ-secretase inhibitors [Inhibitor IV (Calbiochem) and L685,458 (Tocris)] were performed under similar conditions for 12 h. Supernatant was collected and spun down for 10 min at 15000 rpm, meanwhile cells were lysed. 1 mL of supernatant and 200 µl of lysate were incubated with 1.5 µg 6E10 antibody (Biolegend) in 500 µl PBS and 40 µl Protein G sepharose at 4°C rotating overnight. Samples were washed 3x with PBS and using an 18-gauge needle, beads were air dried. Finally, 2x LDS sample buffer with β-mercaptoethanol was added and samples were boiled for 5 min at 95°C. The remaining cell culture supernatant and lysate were prepared for SDS-PAGE as input. For mouse primary cortical cultures, immunoprecipitation was conducted for the supernantant as described above using 2 µg of M3.2 antibody (Biolegend) and 50 µl Protein G sepharose.

### Western blot analysis

Samples were separated on 4-12% bis-tris gels (Novex, Nupage, Invitrogen), 8% tris-glycine or 10-20% tris-tricine (Novex, Nupage, Invitrogen) SDS-PAGE gels for optimal protein and fragment detection. Bis-tris gels were run with MES running buffer (Invitrogen). Proteins were transferred onto nitrocellulose using transfer buffer with 10% ethanol or 20% ethanol for Aβ fragments. For detection of Aβ or Aη fragments, western blots were boiled for 5 min in PBS. For endogenous mouse Aη, Western blots were blocked with 0.2% Tropix i-Block in TBS-T whereas other westerns were blocked with 5% skim milk in TBS-T. The following primary antibodies were used: 22C11 (Millipore), 6E10 (human Aβ region-specific, Biolegend), 2E9 (epitope as determined by Willem et al.: PWHSFGADSVP, N-terminal of β-cleavage and C-terminal of η-cleavage site; Millipore), M3.2 (mouse Aβ region-specific, Biolegend), Y188 (APP C-terminus, ab32136, Abcam), C1/6.1 (APP C-terminus, Biolegend) mouse-anti-myc (9B11, Cell Signaling), mouse-anti-β-actin (8H10D10, Cell Signaling), rabbit-anti-β-tubulin (2128, Cell Signaling), rabbit-anti-flag (D6W5B, Cell Signaling), rabbit-anti- RHBDL4 (HPA013972 Sigma). Horseradish peroxidase (HRP)-coupled secondary antibodies directed against mouse or rabbit IgG were purchased from Promega. Chemiluminescence images were acquired using the ImageQuant LAS 500 or 600 system (GE Healthcare).

### Mass spectrometry

For determining APP cleavage sites, HEK293T cells were transfected with APP containing an N-terminal myc-tag. 36 h post transfection, cells were lysed, 200 µl lysate was incubated with 20 µl anti-myc-sepharose (3400S, Cell signaling) in 200 µl PBS at 4°C over night. Samples were washed twice with lysis buffer, once with PBS + 400 mM NaCl, once with PBS, and then twice with 50 mM ammonium acetate. Elution was performed with 2x300 µl of 50% acetic acid. Eluates were spun down for 10 min at 20,000 x g and then 550 µl of the supernatant was dried in a SpeedVac concentrator (Savant). Samples were digested using Endoproteinase LysC as follows: resuspension in denaturing buffer (6 M urea, 1 mM EDTA, 50 mM Tetraethylammonium bicarbonate (TEAB) pH 8.5), reduction with 2 mM Tris(2- carboxyethyl)phosphine hydrochloride (TCEP) for 10 min at 37°C, blocking of cysteine residues for 30 min in the dark by adding 10 mM iodoacetamide and overnight digestion with 0.25 µg LysC. Digests were purified with C18 ZipTips (Merck) and dried. Samples were dissolved in 0.1% Trifluoroacetic acid/4% acetonitrile, captured on a C18 µ-precolumn (Waters) and eluted onto an Acclaim PepMap100 C18 column (75 µm × 15 cm, Pierce) with a 1 h 5-40% gradient of acetonitrile in 0.1% formic acid at 300 nL/min. The eluted peptides were analyzed with an Impact II Q-TOF spectrometer equipped with a CaptiveSpray electrospray source with an acetonitrile-enriched NanoBooster gas (Bruker, US). Data was acquired using data-dependent auto-MS/MS with a range 150-2200 m/z range, a fixed cycle time of 3 sec, a dynamic exclusion of 1 min, m/z-dependent isolation window (1.5-5 Th) and collision energy in the range 25-75 eV (Beck *et al*, 2015). The raw data was processed using Andromeda, integrated into MaxQuant (Cox *et al*, 2011). A “semi-specific” search with LysC was performed, using the entire human proteome sequence database (Uniprot) and common contaminants provided by MaxQuant. Peptides with non-lysine-residues at the C-terminus and scores above 50 were selected for analysis. Extracted ion chromatograms for selected peptides were integrated using the Data Analysis software (Bruker).

### Mouse brain tissue analysis

For the analysis of Aη-like fragments, we strictly followed the protocol published by Willem et al. (Willem et al., 2015). In brief, approximately 150 mg brain tissue (comparable frontal cortical brain areas) were homogenised with 1:4 weigth per volume diethylamine (DEA) buffer (50 mM NaCl, 0.2% diethylamine, pH 10, cOmplete Protease InhibitorsTM, Roche). After 60 min at 130000xg and 4°C, the DEA fraction was collected and adjusted to pH 6.8. Lysates were adjusted to 5 μg/μl protein concentration and 2x LDS sample buffer with β-mercaptoethanol was added as preparation for SDS-PAGE. Proteins were separated on 10-20% tris-tricine gels. For APP expression in RHBDL4 knockout animals (brain tissue obtained from Dr. Matthew Freeman’s lab, Project license number PP5666180, University of Oxford, UK), similar frontal cortical brain regions were snap frozen after dissection and stored at −80°C until further analysis. For tissue lysis, 5 times the volume of lysis buffer (20 mM Hepes, 150 mM NaCl, 10% glycerol, 2 mM EDTA, 1% NP-40, 0.1 % sodium deoxycholate, pH 7.4) was added to 100 mg of brain tissue. After mechanical tissue homogenization, lysates were incubated at 4°C for 1 h and then spun down at 15000g for an additional hour at 4°C. Lysates were adjusted to equal protein concentrations (3-5 µg/µl) and 2x LDS sample buffer with β- mercaptoethanol was added as preparation for SDS-PAGE.

### RHBDL4 KO mice

Homozygous RHBDL4-knockout mice are viable and show no obvious phenotypes (Lastun *et al*., 2022).Cortical dissociation was conducted on RHBDL4 KO mice originally obtained from Dr. Matthew Freeman’s lab and housed in the Munter lab colony according to the McGill University standard operating procedure mouse breeding colony management #608. All procedures were approved by McGill’s Animal Care Committee and performed in accordance with the ARRIVE guidelines (Animal Research: Reporting in Vivo Experiments). RHBDL4 knockout mice were generated through a deletion of Exon 2 (Lastun *et al*., 2022) and are viable and fertile. Homozygous breeders were maintained, and offspring compared to litters received through WT x HET breeding of the parents’ siblings. Genotyping was performed by Transnetyx.

### Data analysis and statistics

Western blot images were quantified with ImageJ. Statistical data analysis was performed with GraphPad Prism 9. Details as indicated in the figure legends.

## Supplemental information

**Suppl. Figure 1: Sequence validation of identified peptides.**

**A-C)** MS/MS data for three peptides generated by LysC proteolysis of myc-APP-flag immunoprecipitated with an anti-myc antibody. Shown are the b- and y-ion series covering completely **(A)** or almost the entire peptide sequence **(B, C)**. From these data, peptide sequences were identified with Maxquant by matching to a database of human proteins (Uniprot) that were semi-specifically digested with LysC. Data analysis is summarized in the table **(D)**. The mass error was calculated between the predicted and observed precursor mass. The posterior error probability (PEP) is the probability of the peptide being wrongly identified, based on target-decoy analysis. The Maxquant score is a probability based score combining mass error, charge, MS/MS match, etc.

## Acknowledgements

We thank Dr. Michael Willem for initially providing the 2E9 antibody. We thank Meijuan Niu for valuable help with DNA cloning and Felix Oestereich for help in sample preparation for MS. This research was supported by NSERC Discovery grant no. RGPIN-2015-04645, the Canadian Institutes of Health Research (CIHR) PJT-175306 and PJT-162302, Canada Foundation of Innovation Leaders Opportunity Fund (CFI-LOF, 32565), Alzheimer Society of Canada Young Investigator award PT-58872 and Research Grant 17-02, Fonds d’innovation Pfizer-FRQS sur la maladie d’Alzheimer et les maladies apparentées no. 31288, McGill Faculty of Medicine Bridge funding, and The Scottish Rite Charitable Foundation of Canada 16112. YP received the Davis fellowship through McGill’s Faculty of Medicine and a stipend from the Canada First Research Excellence Fund, awarded to McGill University for the Healthy Brains for Healthy Lives initiative. SR received a CGS-Master’s award.

## Author contributions

YP, SP and SJR designed and conducted experiments, analyses and manuscript preparation. CL collected the RHBDL4-knockout mouse brains. FC prepared dissociated mixed cortical cultures. AD prepared samples for mass spectrometry and performed analysis. MF generated RHBDL4-knockout mouse strain. AM supervised experiments with dissociated cultures. JFT analysed mass spectrometry data, LMM supervised and coordinated the project, revised manuscript. All authors revised and approved the manuscript.

