## Supplemental figure for "Eta-secretase-like processing of the amyloid precursor protein (APP) by RHBDL4"

**Peptide 1** **Peptide 2** **Peptide 3**

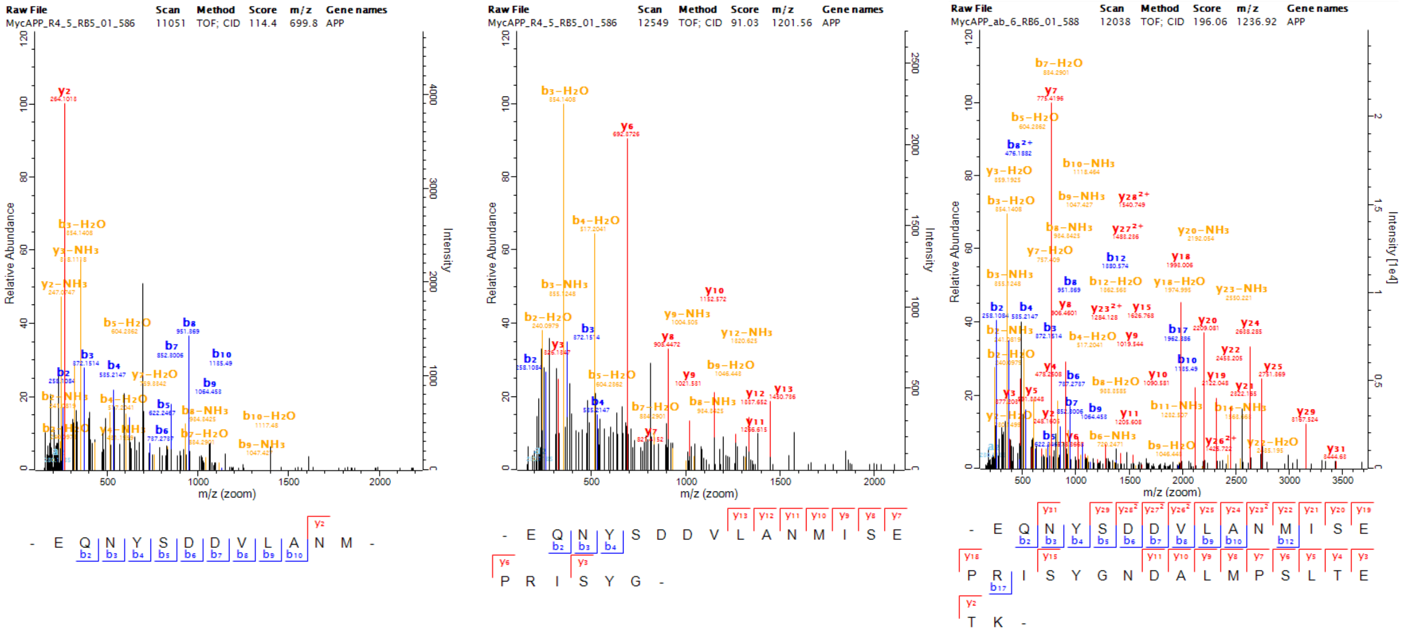


| Peptide # | Sequence | Length | Mass | Charges | P-value | Maxquant score |
|---|---|---|---|---|---|---|
| 1 | EQNYSDDVLANM | 12 | 1397.5769 | 2 | 9.26E-165 | 114.4 |
| 2 | EQNYSDDVLANMISEPRISYG | 21 | 2400.0904 | 2 | 1.10E-41 | 91.032 |
| 3 | EQNYSDDVLANMISEPRISYGNDALMPSLTETK | 33 | 3700.7237 | 3;4 | 0 | 244.26 |

Supplemental Figure 1
